# Trophic shift and the origin of birds

**DOI:** 10.1101/2020.08.18.256131

**Authors:** Yonghua Wu

## Abstract

Birds are characterized by evolutionary specializations of both locomotion (e.g., flapping flight) and digestive system (toothless, crop, and gizzard), while the potential selection pressures responsible for these evolutionary specializations remain unclear. Here we used a recently developed molecular phyloecological method to reconstruct the diets of the ancestral archosaur and of the common ancestor of living birds (CALB). Our results showed that the ancestral archosaur exhibited a predominant Darwinian selection of protein and fat digestion and absorption, whereas the CALB showed a marked enhanced selection of carbohydrate and fat digestion and absorption, suggesting a trophic shift from carnivory to herbivory (fruit, seed, and/or nut-eater) at the archosaur-to-bird transition. The evolutionary shift of the CALB to herbivory may have essentially made them become a low-level consumer and, consequently, subject to relatively high predation risk from potential predators such as gliding maniraptorans, from which birds descended. Under the relatively high predation pressure, ancestral birds with gliding capability may have then evolved not only flapping flight as a possible anti-predator strategy against gliding predatory maniraptorans but also the specialized digestive system as an evolutionary tradeoff of maximizing foraging efficiency and minimizing predation risk. Our results suggest that the powered flight and specialized digestive system of birds may have evolved as a result of their tropic shift-associated predation pressure.

## Introduction

Diet plays a fundamental role in the life of an animal. It defines interactions with other organisms and shapes their evolution. Modern birds exhibit diverse diet preferences, including herbivory, omnivory, and carnivory, while the diet of ancestral birds remains less clear. Fossil evidence shows that ever since the origin of birds from the Late Jurassic, they underwent adaptive radiation to diversified dietary niches in the Cretaceous, with herbivorous (e.g., fruits and seeds), piscivorous, and insectivorous diets found (Zhou and Zhang 2002; Zhou et al. 2003; Benton 2015; Chatterjee 2015; Chiappe and Qingjin 2016; Ksepka et al. 2019; O’Connor 2019; O’Connor and Zhou 2020). In particular, seed and/or fruit eating are suggested in many ancestral bird lineages, such as *Jeholornis*, *Sapeornis*, *Hongshanornis*, and *Yanornis* (Zhou and Zhang 2002; Zheng et al. 2011; Benton 2015; Chatterjee 2015; Chiappe and Qingjin 2016; Mayr 2017; Ksepka et al. 2019; O’Connor 2019; O’Connor and Zhou 2020). This suggests that seed and/or fruit eating may have been relatively common during the early evolution of birds and that this herbivorous adaptation may play a vital role in the early evolution of birds (Zanno and Makovicky 2011; Zheng et al. 2011; Chatterjee 2015; Wang et al. 2016a).

Studies in comparative digestive physiology provide important insights into understanding the molecular bases underlying the dietary variation of animals (Karasov et al. 2011; Karasov and Douglas 2013). Accumulating evidence has revealed a fundamental pattern that the digestive physiology of animals evolves in parallel with their diets (Karasov et al. 2011; Karasov and Douglas 2013; Miller and Harley 2016). And the digestion and absorption capability of animals generally reflects their dietary load of different nutrient substrates such as carbohydrates, proteins, and fats (Corring 1980; Karasov and Diamond 1988; Hidalgo et al. 1998; German et al. 2004; Karasov et al. 2011). The higher the nutrient substrate, the higher the expression and activity of its corresponding digestive enzymes and nutrient transporters and vice versa (Karasov et al. 2011; Karasov and Douglas 2013). This suggests that the digestion and absorption capability of animals is under evolutionary adaptation to approximately match loads of different dietary components such as carbohydrates, proteins and fats in their diets (Schondube et al. 2001; Karasov et al. 2011; Karasov and Douglas 2013; Wang et al. 2016c; Hecker et al. 2019; Wang et al. 2020). With this in mind, one would expect that herbivores and carnivores would tend to present an evolutionary enhancement of the digestion and absorption of plant food and meat, respectively. Regarding plant food and meat (including both invertebrates and vertebrates), one of the important differences between them is that meat is generally high in proteins and fats, while plant food is generally high in carbohydrates (Clinic 2002; Karasov et al. 2011; Chen and Zhao 2019; Hecker et al. 2019; Wang et al. 2020) except seeds, particularly nuts, which are rich in fat as well (Clinic 2002). Indeed, recent studies on the evolution of digestive system-related genes have shown that carnivores more likely show an evolutionary enhancement of the genes related to the digestion and absorption of proteins and fats, while animals consuming abundant plant foods (e.g., herbivores and omnivores) tend to exhibit an evolutionary enhancement of the genes related to the digestion and absorption of carbohydrates (Chen and Zhao 2019; Wang et al. 2020; Wu et al. 2020), with the exception of parrots, which ingest seeds and nuts and present an evolutionary enhancement of the digestion and absorption of fat in addition to carbohydrates (Wu et al. 2020). This may suggest that the adaptive evolution of digestive system-related genes is capable of reflecting the dietary variations of different animals (Chen and Zhao 2019; Wang et al. 2020; Wu et al. 2020).

The recent development of a molecular phyloecological approach, which employs the phylogenetic evolutionary analyses of the molecular markers indicative of trait states, allows us to reconstruct ancestral traits using molecular data (Wu et al. 2017; Wu et al. 2018; Wu and Wang 2019). The substantial dietary differences between carnivores and herbivores in terms of the amounts of dietary components (e.g., carbohydrates, proteins, and fats) (Clinic 2002; Karasov et al. 2011; Hecker et al. 2019; Wang et al. 2020), and the adaptation of digestive system-related genes to the variations of dietary components of animals (Schondube et al. 2001; Karasov et al. 2011; Karasov and Douglas 2013; Wang et al. 2016c; Hecker et al. 2019; Wang et al. 2020; Wu et al. 2020) may suggest that digestive system-related genes can be used as the molecular markers of diets to reconstruct the diets of ancestral animals in the context of molecular phyloecology (Wu et al. 2020). In this study, we employed the molecular phyloecological approach using digestive system-related genes as molecular markers to infer the diets of the ancestral archosaur and of the common ancestor of living birds (CALB). Our results revealed a diet shift from carnivory to herbivory at the archosaur-to-bird transition. The molecular findings of the diet shift, coupled with the research advance of avian paleontology, provide new insights into understanding the origin of birds.

## Results

We examined the adaptive evolution of 83 digestive system-related genes (Table S1) in the context of sauropsid phylogeny (Fig. 1). The 83 genes came from the three pathways related to carbohydrate digestion and absorption (CDA), protein digestion and absorption (PDA), and fat digestion and absorption (FDA) (Fig. 2). Following the molecular phyloecological approach to reconstruct ancestral traits (Wu et al. 2017; Wu et al. 2018; Wu and Wang 2019), we used branch and branch-site models implemented in PAML software (Yang 2007) to detect positively selected genes (PSGs) along our target branches, and PSGs were found based on branch-site model (Tables S2-3). The evidence of the positive selection of genes may suggest a functional enhancement of their corresponding functions in relevant lineages (Wu et al. 2017; Wu et al. 2018; Wu and Wang 2019).

**Fig.1.**
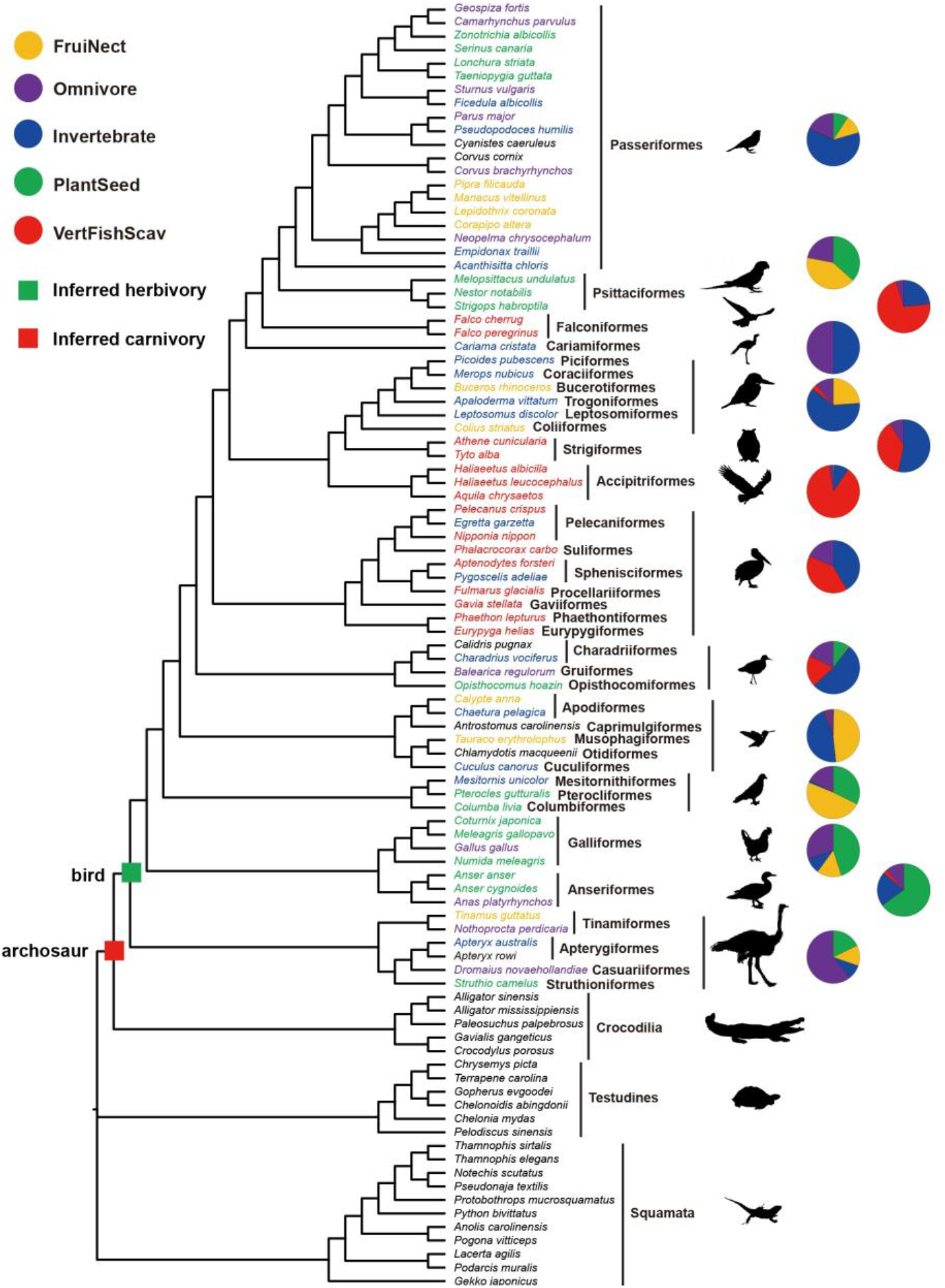
Phylogeny and the diets of modern birds. Phylogenetic relationships of taxa used follow published studies (Jønsson and Fjeldså 2006; McKay et al. 2010; Oaks 2011; Guillon et al. 2012; Pyron et al. 2013; Crawford et al. 2015; Wu and Wang 2019) and the Tree of Life Web Project (http://tolweb.org/tree/). Dietary categories of each bird species follow one published study (Wilman et al. 2014) and are shown in different colors, and the bird species without dietary information available are shown in black. The dietary categories of avian clades based on the dietary data of a total of 9993 extant bird species are shown in pet charts. PlantSeed (plant and seeds), FruiNect (fruits and nectar), Invertebrate (invertebrates), VertFishScav (vertebrates and fish and carrion) and Omnivore (score of <= 50 in all four categories).

**Fig.2.**
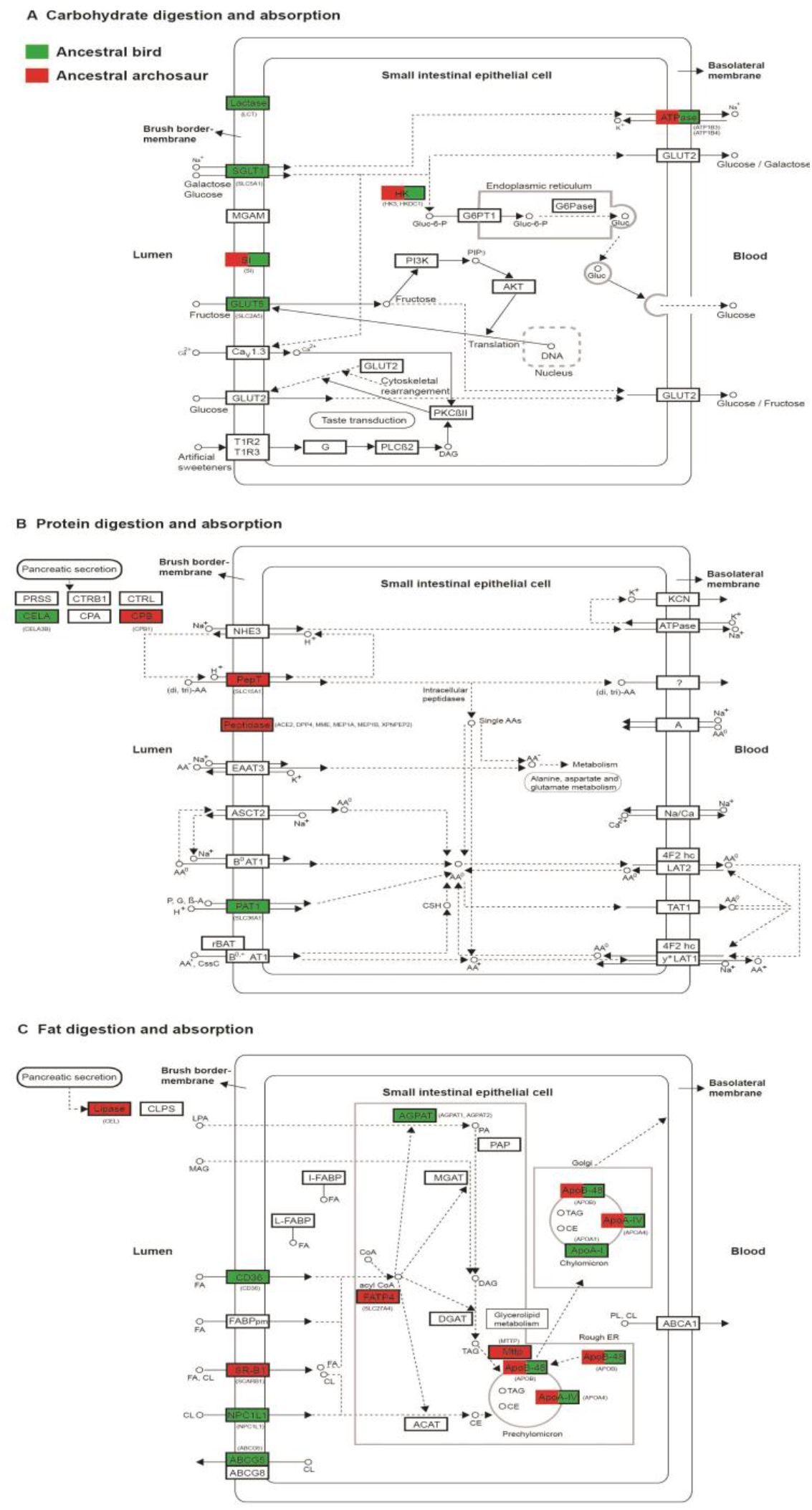
Digestive system pathways and positively selected genes. The digestion and absorption of carbohydrates (A), proteins (B) and fats (C) are shown. The proteins with their corresponding genes in parentheses under positive selection are highlighted in red (ancestral archosaur) and green (ancestral bird). The three digestive system pathways are modified from corresponding KEGG pathways with accession numbers (map04973, map04974, and map04975).

We initially analyzed the positive selection of the digestive system-related genes along the common ancestor branch of living birds. Among the 83 genes analyzed, we found 17 PSGs across all three pathways, with CDA and FDA showing the most predominant positive selection, and PDA showing the relatively weakest positive selection in terms of p values and the number of PSGs (Table S2, Fig. 2). For CDA, seven PSGs (*ATP1B3*, *ATP1B4*, *HK3*, *SLC5A1*, *LCT*, *SI*, and *SLC2A5*) were found with generally low p values compared to those PSGs found in FDA and PDA. Among the seven PSGs found, three genes, *SI*, *LCT*, and *HK3*, showed the most predominant positive selection signals. *SI* encodes sucrase-isomerase and is essential for the digestion of dietary carbohydrates, such as starch, sucrose, and isomaltose (Naim et al. 1988). *LCT* shows lactase activity and phlorizin hydrolase activity (Boll et al. 1991). *HK3* is involved in glucose metabolism (Furuta et al. 1996). Similar to *HK3*, one positively selected gene, *SLC5A1*, also plays a role in glucose metabolism, functioning as a transporter of glucose in the small intestine (Wright et al. 2007). Intriguingly, we detected the positive selection of one gene *SLC2A5*, encoding GLUT5, which is known to have an exclusive affinity for fructose (fruit sugar) and is the major fructose transporter in the intestines and other tissues, mediating the uptake of dietary fructose (Douard and Ferraris 2008; Cura and Carruthers 2012; Mueckler and Thorens 2013). We also detected the positive selection of *ATP1B3* and *ATP1B4*, which encode the Na^+^/K^+^ ATPase involved in the CDA pathway to maintain ionic homeostasis (Li et al. 2011). Besides CDA, we detected PSGs involved in the FDA pathway, and eight PSGs (*ABCG5*, *AGPAT1*, *AGPAT2*, *APOA1*, *APOA4*, *APOB*, *CD36* and *NPC1L1*) were found. Of these, *CD36* plays an important role in the uptake and processing of fatty acids (Pepino et al. 2014). *NPC1L1* is involved in the intestinal absorption of cholesterol and plant sterols (Izar et al. 2011). *AGPAT1* and *AGPAT2* play roles in converting lysophosphatidic acid (LPA) into phosphatidic acid (PA) (Takeuchi and Reue 2009). *APOA1*, *APOA4*, and *APOB* encode key apolipoproteins to carry fats and fat-like substances in the blood (Mangaraj et al. 2016; Qu et al. 2019). Remarkably, we found the positive selection of one gene, *ABCG5*, which encodes sterolin-1 and works together with sterolin-2, encoded by gene *ABCG8*, to form a protein called sterolin. Sterolin is a transporter protein and plays an important role in eliminating plant sterols to regulate the whole-body retention of plant sterols (Hazard and Patel 2007; Izar et al. 2011), which are mainly present in nuts and seeds (Izar et al. 2011). Unlike CDA and FDA, relatively weak positive selection signals were found in PDA with only two PSGs (*CELA3B* and *SLC36A1*) detected, which are respectively involved in digesting dietary proteins and transporting intestinal amino acids (Frølund et al. 2010; Szabó et al. 2016).

We subsequently examined the positive selection of the digestive system- related genes along the branch leading to the common ancestor of living birds and crocodilians, representing the ancestral archosaur. In contrast to the CALB, for the ancestral archosaur, we detected the highest number of PSGs in PDA, followed by FDA, with the lowest number of PSGs found in CDA (Fig. 2, Table S3). For PDA, eight PSGs were found, of which seven PSGs (*ACE2*, *CPB1*, *DPP4*, *MEP1A*, *MEP1B*, *MME*, and *XPNPEP2*) encode peptidases (Yamamoto et al. 1992; Tipnis et al. 2000; Lambeir et al. 2003; Crisman et al. 2004; Erşahin et al. 2005; Higuchi et al. 2016), and one gene, *SLC15A1*, is involved in the intestinal transport of peptide (Liang et al. 1995). For FDA, we found six PSGs, including *APOA4*, *APOB*, *CEL*, *MTTP*, *SCARB1*, and *SLC27A4*. Of which, *CEL* encodes a bile salt-dependent carboxyl-ester lipase, hydrolyzing dietary fat, and cholesteryl esters in the small intestine (Johansson et al. 2018). *SCARB1* mediates the uptake of cholesterol and lipids (Shen et al. 2018). *SLC27A4* is an important fatty acid transporter in small intestinal enterocytes (Stahl et al. 1999). *MTTP* is involved in the transport of triglycerides, cholesteryl esters, and phospholipids (Hussain et al. 2012). *APOA4* and *APOB* encode two key apolipoproteins responsive to carrying fats and fat-like substances in the blood (Mangaraj et al. 2016; Qu et al. 2019). Unlike PDA and FDA, CDA showed the lowest number of PSGs, and only three PSGs (*SI*, *HKDC1*, and *ATP1B4*) were detected. *HKDC1* is involved in glucose metabolism (Ludvik et al. 2016). *ATP1B4* encodes Na^+^/K^+^ ATPase to maintain ionic homeostasis (Li et al. 2011). *SI* encodes sucrase-isomerase, digesting dietary carbohydrates including starch, sucrose, and isomaltose (Naim et al. 1988).

Our positive selection analyses described above demonstrated that the CALB showed a predominant Darwinian selection of the genes related to carbohydrate and fat digestion and absorption, whereas the ancestral archosaur exhibited a marked Darwinian selection of the genes related to protein and fat digestion and absorption, indicating substantial selection differences between them (Fig. 2, Tables S2-3). To further know the possible selection differences between the CALB and the ancestral archosaur, we then used the program RELAX (Wertheim et al. 2015) to examine the selection intensity changes of the digestive system-related genes of the CALB relative to those of the ancestral archosaur (Table S4). Our results showed that FDA-related genes exhibited the most intensified selection, followed by CDA-related genes, while PDA-related genes showed the relatively weakest selection intensification. For FDA, eight genes (*ABCA1*, *AGPAT1*, *CD36*, *FABP1*, *MTTP*, *NPC1L1*, *GOT2*, and *SCARB1*) exhibited a relative intensified selection. Among the eight genes, five (*AGPAT1*, *CD36*, *MTTP*, *NPC1L1*, and *SCARB1*), as mentioned above, are mainly related to the transport or conversion of lipids, while the other three genes, *ABCA1*, *FABP1*, and *GOT2*, are mainly involved in the transport of lipids. Specifically, *ABCA1* mediates cellular cholesterol and phospholipid efflux (Wang and Tall 2003), *FABP1* regulates lipid transport and metabolism (Wang et al. 2015), and *GOT2* is involved in fatty acid transport (Tousignant et al. 2019). Besides FDA, three CDA-related genes, *PIK3CB*, *SLC37A4*, and *CACNA1D*, showed selection intensification as well. *PIK3CB* encodes the catalytic subunit of phosphoinositide 3-kinase (PI3K), which plays a role in regulating the activity of GLUT5, a major fructose transporter (Cui et al. 2005). *SLC37A4* acts as a transporter of glucose 6-phosphate (Cappello et al. 2018). *CACNA1D* encodes a subunit of a calcium channel (CaV1.3) (Fig. 2). For PDA, only one gene, *CELA3B*, which encodes pancreatic serine proteinases (Szabó et al. 2016), was subject to selection intensification. In addition to these selection-intensified genes, several genes showed relative selection relaxation in the CALB compared to the ancestral archosaur, including two genes (*ABCG5* and *APOB*) of FDA, one gene (*CTRL*) of PDA and one gene (*SLC5A1*) clearly involved in CDA.

## Discussion

### Diet shift (carnivory to herbivory) at the archosaur-to-bird transition

Comparative digestive physiology studies demonstrate that the evolution of digestive system molecules adapts for the amounts of nutrient components (e.g., carbohydrates, fats and proteins) in the diets of animals (Karasov et al. 2011; Karasov and Douglas 2013). Our results showed that the ancestral archosaur exhibited a marked selection of the genes related to protein and fat digestion and absorption, while the CALB presented a predominant selection of the genes involved in carbohydrate and fat digestion and absorption (Fig. 2, Table S2-4). Especially for the ancestral archosaur, our positive selection analyses revealed the highest number of PSGs in PDA, followed by FDA, with the lowest number of PSGs found in CDA (Fig. 2). This may suggest that the diet of the ancestral archosaur was characterized by a high amount of proteins, followed by fats, with a minimum load of carbohydrates. This nutrient profile is highly consistent with the presumable carnivory of ancestral archosaurs (Nesbitt 2011) since meats are generally rich in proteins, followed by fats, with the minimum amount of carbohydrates (Clinic 2002). Contrary to the ancestral archosaur, for the CALB, our results based on two different methods (PAML and RELAX) consistently demonstrated that it showed the most predominant selection in carbohydrate and fat digestion and absorption-related genes, with the weakest selection found in PDA-related genes (Fig. 2, Table S2). This may suggest that the diet of the CALB is characterized by high amounts of carbohydrates and fats, with a relatively minimal amount of proteins, representing a high-energy diet. This seems to be more consistent with herbivory compared to carnivory, considering that plant foods are rich in carbohydrates, whereas meats are particularly high in proteins (Clinic 2002). In particular, most PSGs involved in CDA (*SI*, *SLC5A1*, *SLC2A5*, *HK3*, and *LCT*) found in the CALB center on the digestion and absorption of sugars (e.g., glucose, sucrose and fruit sugar), indicating its high-sugar diet. A high-sugar diet may suggest their eating of fruits, which are characterized by relatively high amounts of sugars among plant foods (Schondube et al. 2001; Clinic 2002; Yahia 2018). In particular, one positively selected gene, *SLC2A5*, found in the CALB is mainly involved in the transports of fruit sugar (Douard and Ferraris 2008; Cura and Carruthers 2012; Mueckler and Thorens 2013). These lines of evidence may suggest that the CALB involved fruits in its diet. On the other hand, for the PSGs found in FDA, one gene, *ABCG5*, plays a critical role in the transport of plant sterols, which are mainly found in nuts and seeds (Hazard and Patel 2007; Izar et al. 2011). This may suggest that the CALB ingests seeds and/or nuts as well, which are rich in fat (Clinic 2002). The predominant selection of the CALB in FDA is similar to parrots, which consume considerable amounts of seeds and nuts and are found to present a strong Darwinian selection in FDA as well with four PSGs found, of which three (*ABCG5*, *APOA4*, and *APOB*) are shared with the CALB (Wu et al. 2020). In all, our molecular study suggests that the ancestral archosaur is probably a carnivore, while the CALB is more likely an herbivore, ingesting fruits, seeds, and/or nuts (Fig. 1).

Our molecular results are highly consistent with the fossil evidence showing that ancestral archosaurs are generally typically meat eaters (Nesbitt 2011), and a great number of ancestral Mesozoic birds, including the basal birds, such as *Jeholornis*, *Confuciusornis* and *Sapeornis*, show features or gut contents indicating that they ate fruits and/or seeds (Zhou and Zhang 2002; Chatterjee 2015; Chiappe and Qingjin 2016; Mayr 2017; Ksepka et al. 2019; O’Connor 2019; O’Connor and Zhou 2020). Particularly, for the herbivory of the CALB found in this study, it is consistent with the widespread herbivory observed in many living bird lineages across bird phylogeny (Fig. 1). In line with this, one previous study shows evidence of seeds as an important dietary component of the CALB using maximum likelihood reconstruction (Larson et al. 2016). Considering that ancestral birds lived in a conifer-dominated ecosystem (Zhou et al. 2003; Matsukawa et al. 2014), the seeds that they ate might partly come from conifers (Zheng et al. 2011). Indeed, the seeds of many conifers (e.g., pines) are relatively rich in lipids (Wolff et al. 2000; Wolff et al. 2002), which might have led to the evolutionary enhancement of FDA of the CALB found in this study. Additionally, previous studies show that the Late Jurassic/Early Cretaceous radiation of more advanced birds temporally coincides with that of angiosperm plants (Friis et al. 2011), and it is likely that the fruit- and/or seed-eating habitat of ancestral birds may have partly helped for their dispersal of seeds (Mayr 2017). The herbivory of the CALB is also consistent with the occurrence of ceca observed in the majority of living birds, including the basal lineages (e.g., ratites), which is generally considered to be helpful for cellulose digestion and fermentation linked to herbivory (Clench and Mathias 1995; Miller and Harley 2016). The dietary shift of the CALB to herbivory is also consistent with the observation of reductions in both teeth and biting force across the theropod-bird transition, which is considered to have resulted from the dietary shift from carnivorous to herbivorous diets (Zanno and Makovicky 2011; Li et al. 2020). The similar transition from carnivory to herbivory occurs multiple times in theropods (Zanno et al. 2009; Zanno and Makovicky 2011; Barrett 2014). The causes underlying the evolutionary shift to the herbivory of the CALB are not clear, but the possible competition from carnivorous theropods and pterosaurs is proposed as a possible candidate (Wang et al. 2016a; Li et al. 2020). The finding of the herbivory of the CALB ingesting fruits, seeds, and/or nuts, which characterize seed plants adapted to dry land environments (Cowen 2013), may strongly suggest that the CALB mainly occurred in terrestrial habitats rather than an aquatic environment, as hypothesized previously (You et al. 2006). These findings are consistent with the fact that the phylogenetically most basal extant neornithine birds—that is, Palaeognathae and Galloanseres—are predominantly herbivorous or omnivorous and mainly occur in terrestrial habitats (Mayr 2017).

### Trophic shift of ancestral birds as low-level consumers

Our results suggest an evolutionary shift of the CALB to an herbivorous diet (fruit, seed and/or nut eater) (Fig. 1). Evolutionarily, birds are widely believed to be derived from a group of small maniraptoran theropods, including dromaeosaurids and troodontids (Xu et al. 2014; Benton 2015; Chatterjee 2015; Mayr 2017). Among these maniraptoran theropods, many of them, including most dromaeosaurids and derived troodontids, show carnivory (Zanno et al. 2009; O’Connor et al. 2011; Xu et al. 2011; Zanno and Makovicky 2011; Brusatte 2012; Han et al. 2014; Chatterjee 2015; Button et al. 2017; Mayr 2017; O’Connor 2019; O’Connor et al. 2019; O’Connor and Zhou 2020) (Fig. 3). However, unlike their maniraptoran relatives, many bird lineages, including the basal bird lineages, such as *Jeholornis* and *Sapeornis*, may have evolved to exploit herbivorous niches, as evidenced by both the molecular (Fig. 1) and fossil evidence mentioned above (Zhou and Zhang 2002; Zanno and Makovicky 2011; Zheng et al. 2011; Benton 2015; Chatterjee 2015; Chiappe and Qingjin 2016; Mayr 2017; Ksepka et al. 2019; O’Connor 2019; Li et al. 2020; O’Connor and Zhou 2020) (Fig. 3). The dietary shift from carnivory to herbivory may suggest a shift of the trophic niche of bird ancestors from that of a high-level consumer to a low-level consumer as a primary and/or secondary consumer (Matsukawa et al. 2014; O’Connor et al. 2019). This is consistent with the marked reduction or loss of teeth along with the evolution of birds (Benton 2015; Chatterjee 2015), a feature indicative of low-level consumers rather than high-level consumers (e.g., top predators), which would otherwise show a predation feature of well-developed teeth (Farlow and Holtz 2002; Brusatte 2012). Moreover, though diverse diets (e.g., seeds, fish, and insects) among ancestral bird lineages have been found, there is no direct fossil evidence indicative of their preying on terrestrial vertebrates (O’Connor 2019), strengthening their ecological niches as low-level consumers. Ancestral birds were abundant in Mesozoic terrestrial ecosystems (Matsukawa et al. 2014), occurring globally and representing a potential food source for carnivores. Becoming a low-level consumer, ancestral birds may be under increased predation risk. This is particularly the case for ancestral birds since they evolve toward miniaturization suitable for powered flight (Benson et al. 2014; Lee et al. 2014), and their small body size may be particularly vulnerable to predators. More importantly, their evolution of endothermy and powered flight requires much more energy and, consequently, frequent foraging (Chatterjee 2015; O’Connor and Zhou 2015; O’Connor 2019; Wu and Wang 2019). Frequent foraging may have most often exposed them to predators, hence leading to their high predation risk. In support of this, fossil evidence shows that ancestral birds, such as enantiornithines and *Confuciusornis*, have a precocial development style (Chatterjee 2015; Chiappe and Qingjin 2016), which is generally considered to be an anti-predation strategy for facing historically strong predation pressure (Arendt 1997; Jackson et al. 2009). Moreover, one recent study shows that the CALB was probably cathemeral (i.e., active in both day and night) and that it may have evolved an enhanced visual capability to detect motion (Wu and Wang 2019). Cathemerality is considered to be linked to high predation risk (Colquhoun 2006; Maor et al. 2017), and the promoted motion detection ability of the CALB may mainly help to detect approaching predators given its herbivory. Therefore, the dietary shift may have made ancestral birds become the prey of high-level consumers, possibly leading to their high predation risk.

**Fig.3.**
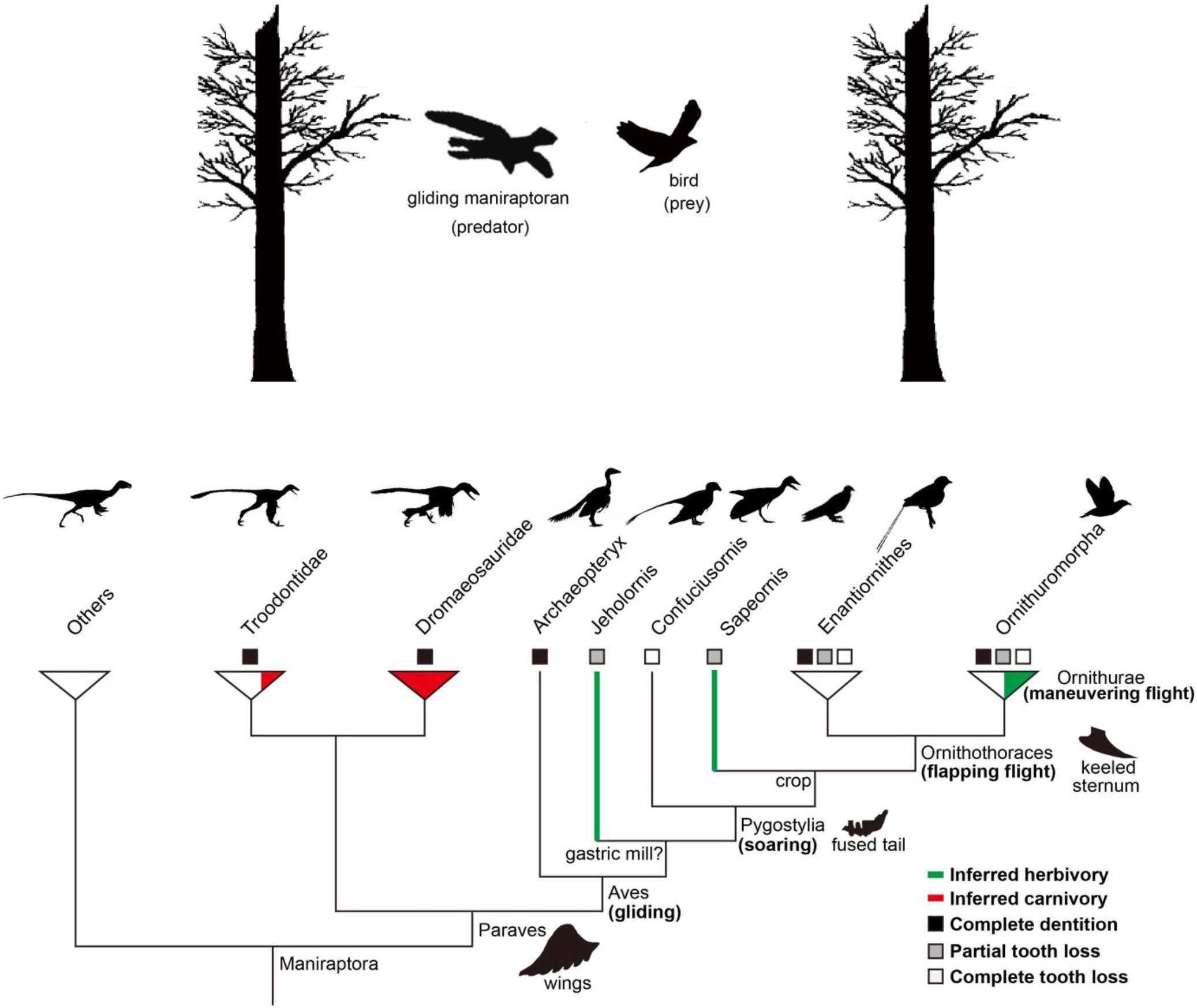
Schematic representation of the predation hypothesis underlying the origin of birds proposed in this study. The predation of gliding predatory maniraptorans on ancestral birds in the context of the arboreal theory is shown (please see text for details). Maniraptoran phylogeny with digestive system characteristics (gastric mill, crop, and tooth) is based on one previous study (O’Connor 2019). The dietary information follows published studies (Zanno et al. 2009; Zanno and Makovicky 2011; O’Connor 2019). The silhouettes of flight-related anatomical features (wings, fused tail, and keeled sternum) along phylogeny are modified from one published study (Brusatte et al. 2015). The progressive enhancement of flight performance from gliding to soaring, flapping flight and maneuvering flight within Aves is based on published literature (Chatterjee 2015). Species silhouettes corresponding to each of phylogenetic taxa used are from phylopic.org and designed by (from left to right): Others (Maniraptora) (Hanyong Pu, Yoshitsugu Kobayashi, Junchang Lü, Li Xu, Yanhua Wu, Huali Chang, Jiming Zhang, Songhai Jia & T. Michael Keesey), Troodontidae (Scott Hartman), Dromaeosauridae (Scott Hartman (modified by T. Michael Keesey)), *Archaeopteryx* (Dann Pigdon), *Jeholornis* (Matt Martyniuk), *Confuciusornis* (Scott Hartman), *Sapeornis* (Matt Martyniuk), Enantiornithes (Matt Martyniuk) and Ornithuromorpha (Juan Carlos Jerí). The tree silhouettes at the top of the image are modified based on published literature (Chatterjee 2015).

### Main predators of ancestral birds—gliding maniraptorans

Arboreal theory proposes that birds evolved from a group of arboreal and gliding maniraptorans and that ancestral birds may be primarily arboreal and capable of gliding flight though they spent some time on the ground as well (Xu et al. 2000; Zhou and Zhang 2002; Xu et al. 2003; Benton 2015; Chatterjee 2015; Chiappe and Qingjin 2016). Given the possible arboreality and gliding lifestyle of ancestral birds, while there are many potential predators, such as carnivorous theropods, carnivorous mammals (e.g., *Repenomamus*), snakes (e.g., *Sanajeh*) and crocodylomorphs, in the Mesozoic terrestrial ecosystem (Brusatte 2012; Matsukawa et al. 2014), four lines of evidence may suggest that one group of carnivorous theropods—maniraptorans (e.g., dromaeosaurids)—is likely one of the main predators of ancestral birds, as proposed previously (Gong et al. 2010; O’Connor et al. 2011; Chatterjee 2015). Primarily, a wealth of small feathered maniraptorans, such as *Aurornis*, *Anchiornis*, *Bambiraptor*, *Buitreraptor*, *Changyuraptor*, *Eosinopteryx*, *Jinfengopteryx*, *Microraptor*, *Rahonavis*, and *Xiaotingia*, are found to have hallmark anatomical characteristics indicative of their capability of gliding flight or even some forms of powered flight (Han et al. 2014; Xu et al. 2014; Benton 2015; Chatterjee 2015; Chiappe and Qingjin 2016; Mayr 2017; Sullivan et al. 2017; Pei et al. 2020), and many of these volant maniraptorans, such as *Microraptor*, *Anchiornis*, and *Changyuraptor*, show predatory features (Zanno et al. 2009; Gong et al. 2010; O’Connor et al. 2011; Xu et al. 2011; Zanno and Makovicky 2011; Brusatte 2012; Han et al. 2014; Button et al. 2017; Mayr 2017; O’Connor 2019; O’Connor et al. 2019; O’Connor and Zhou 2020), representing one of the potential aerial predators of ancestral birds. The predation pressure from these aerial predators may be more significant than those ground predators given the arboreality and gliding lifestyle of ancestral birds.

On the other hand, both ancestral birds and gliding maniraptorans have a relatively small body size among dinosaurs known (Turner et al. 2007; Brusatte 2012; Benson et al. 2014; Lee et al. 2014), suggesting that ancestral birds may be suitable prey for them. This is because there is a general positive correlation of body size between predators and their target prey, and small predators tend to prey on small prey (Gittleman 1985; Vézina 1985; Carbone et al. 1999; Radloff and Du Toit 2004). Moreover, previous studies show that, among theropods, non-avian maniraptorans show a relatively high metabolic level (e.g., endothermy) comparable to birds (Seebacher 2003; Rezende et al. 2020), suggesting that they possibly had a relatively high activity level. The high activity level of maniraptorans supports the feasibility of their predation on ancestral birds. Finally, and more importantly, there is already direct fossil evidence indicative of the predation of ancestral arboreal bird (adult enantiornithine bird) by arboreal and gliding predatory maniraptorans, such as *Microraptor*, which is known from hundreds of specimens, despite the extreme scarcity of preserved fossils (O’Connor et al. 2011). Additionally, the possible predation of ancestral birds by another predatory maniraptoran, *Sinornithosaurus*, which might be capable of gliding flight (Chatterjee and Templin 2004), has also been proposed previously (Gong et al. 2010). These lines of ecological and fossil evidence suggest that the predation pressure of ancestral birds during their early evolution may, at least partly, mainly have come from those arboreal and gliding maniraptorans.

### Flapping flight as a possible anti-predator strategy against gliding maniraptorans

Different theories have been proposed to account for the origin of the powered flight (e.g., flapping flight) of birds (Benton 2015; Chatterjee 2015). And arboreal theory invokes a natural transition of powered flight via gliding flight (Xu et al. 2000; Zhou and Zhang 2002; Xu et al. 2003; Benton 2015; Chatterjee 2015), but a basic question remains: what was the selection pressure for the natural transition (Hedenström 1999)? While gliding flight is common among both living and extinct animals, powered flight is rare and only known in insects, pterosaurs, birds, and bats (Dudley et al. 2007), suggesting that powered flight may less likely occur without certain selection pressures. This is particularly true for birds since their powered flight demands high energy and substantial evolutionary alternation (e.g., keeled sternum and flight muscles) compared to gliding flight, a simple and cheap way of flying (Chatterjee 2015). Early birds, such as *Archaeopteryx* and *Jeholornis*, are believed to be primarily arboreal and capable of gliding flight, which are believed to be descended from maniraptorans that had already evolved gliding flight (Xu et al. 2003; Benton 2015; Chatterjee 2015). Indeed, many maniraptorans possess asymmetric flight feathers to generate lift, and in particular, the discovery of many bird-like paravians, such as *Microraptor*, *Anchiornis*, *Xiaotingia*, and *Aurorornis*, is the most unusual in developing four wings, suggesting their possible high performance of gliding flight (Benton 2015; Chatterjee 2015; Pei et al. 2020). However, given the diet divergence between non-avian maniraptorans and ancestral birds, and particularly that many of gliding maniraptorans (e.g., *Microraptor* and *Sinornithosaurus*) were potential predators of early birds (Gong et al. 2010; O’Connor et al. 2011; Chatterjee 2015), it is plausible that early birds may have then evolved powered flight (e.g., flapping flight) based on their gliding flight to escape from gliding predatory maniraptorans. The predation pressure from gliding predatory maniraptorans may have worked as a driver to stimulate the evolution of powered flight of their arboreal prey. And the flapping flight of birds may be critical to flee from those gliding predators. Fossil evidence shows that ever since the evolutionary divergence of early birds from their maniraptoran relatives, the evolution of birds has shown a major trend in the improvement of flight, such as from gliding to flapping and maneuvering flight with the acquisition of flight-related characteristics such as a shortening of the tail and a keeled sternum (Xu et al. 2014; Benton 2015; Chatterjee 2015; Mayr 2017) (Fig. 3). The continuous evolutionary enhancement of the flight of ancestral birds may essentially help for an increase of speed and maneuverability of locomotion, both of which are considered to be crucial for escape success (Clemente and Wilson 2016). This may be the case particularly for birds since they could not become large in body size, a potential antipredator strategy observed in many animals (Caro 2005), to evade predators due to their miniaturization constraints in favor of flight. Indeed, for many birds, flying is an important means used to escape from predators (Van den Hout et al. 2010; Chatterjee 2015), suggesting predation is an important selection pressure for powered flight (Dudley et al. 2007; Chatterjee 2015). This is consistent with the observation that birds frequently become flightless in predator-free islands (Wright et al. 2016). Possibly, there was some form of coevolution between ancestral birds and those gliding predatory maniraptorans. And the predation pressure from gliding predatory maniraptorans may be an important candidate contributing to the evolutionary shift from gliding flight to powered flight at the theropod-to-bird transition, though it remains unknown as to whether there were gliding predators other than maniraptorans contributing to the evolution of powerful flight of birds as well.

### Specialized digestive system of birds responsive to predation pressures

Living birds are toothless, and they swallow their food whole, which is temporally stored in their crop and then grinded up by their muscular gizzard. Fossil evidence shows that the specialization of the digestive system occurs in multiple lineages of ancestral birds (Zhou et al. 2010; Zheng et al. 2011; Zheng et al. 2014; Chiappe and Qingjin 2016; Wang et al. 2016b; O’Connor 2019; O’Connor and Zhou 2020) (Fig. 3). A recent genomic study shows that modern birds lost their teeth since their common ancestor about 116 million years ago (Meredith et al. 2014). Regarding the evolutionary specialization of the digestive system of birds, its adaptive significance is, however, less clear. Previous studies indicate that the loss of teeth in birds seems to be linked to an herbivorous diet, which is consistent with the herbivory of the CALB found in this study, but the underlying mechanism remains unknown (Zanno and Makovicky 2011; Zheng et al. 2011; Barrett 2014; Chatterjee 2015). Optimal foraging theory states that predation has a profound influence on the foraging strategies of animals, and animals must trade off two conflicting demands of maximizing foraging efficiency and minimizing predation risk (Lima 1985; Lima et al. 1985; Verdolin 2006). In light of this optimal foraging theory, for the evolutionary specialization of the digestive system of birds, we propose here that herbivores (e.g., ancestral birds) are low-level consumers and, consequently, at relatively high risk to predators. Under high predation risk, the time needed to acquire and process food using teeth may be limited, but the evolutionary specialization of the digestive system of birds may allow them to gather more food as fast as possible (maximizing foraging efficiency) as food can be stored in their crop without expending too much time processing it using their teeth, and then they can seek a safe place to process their food via their gizzard (minimizing predation risk). Consequently, the reduced reliance on teeth for the processing of food as a result of predator avoidance may have then led to the selection relaxation of teeth, leading to their subsequent reduction or loss thereof. This may be particularly true for early birds that would necessarily demand frequent foraging (O’Connor 2019) and much time for the oral processing of their food (e.g., hard seeds) (O’Connor and Zhou 2020) if no gizzard were available under relatively high predation risk (including both aerial, arboreal, and ground predators), while the evolution of the bird-like digestive system may help to maximize foraging efficiency and minimize their exposure to predators. Indeed, direct fossil evidence, for instance, the preservation of intact seeds or whole fish in the alimentary canal of ancestral birds, demonstrates that ancestral birds seem not to have used their teeth to process food; rather, their teeth, if any, were mainly used for the acquisition of food (Zhou and Zhang 2002; Zheng et al. 2011; Zheng et al. 2014; O’Connor and Zhou 2015; Li et al. 2020).

Regarding the reduction or loss of the teeth of birds, it is traditionally attributed to lightening the body for flight (Zhou et al. 2010; Zheng et al. 2011). This, however, cannot explain the occurrence of numerous toothed Mesozoic birds (e.g., Enantiornithes and *Ichthyornis*) (Zhou et al. 2010; Yang and Sander 2018; O’Connor 2019), and hence teeth were probably not a limiting factor for flight (O’Connor and Zhou 2015; Chiappe and Qingjin 2016; Mayr 2017; Zhou et al. 2019). Alternatively, teeth reduction or loss is considered to be partly due to the functional replacement by the muscular gizzard (Louchart and Viriot 2011; Zheng et al. 2014; O’Connor 2019). However, this raises a new question: given that teeth and muscular gizzard have a similar function, why the teeth got lost rather than muscular gizzard? One possibility is that it must expend considerable time processing food using teeth without a gizzard during foraging, which may then largely increase their predation risk. In line with this reasoning, the crop is also suggested to help to gather more food quickly to avoid competitors and/or predators (Zheng et al. 2011; Zheng et al. 2014; Miller and Harley 2016), though an alternative explanation exists (O’Connor and Zhou 2015; O’Connor 2019). Given the possible importance of predation, we argue that the evolution of digestive system characteristics of birds, including teeth reduction or loss, crop and gizzard, are not independent; rather, their evolution is probably mutually dependent. The integrative and/or collective evolution of these characteristics may be a result of both maximizing foraging efficiency and minimizing predation risk. Predation pressure is also believed to be a potential selection pressure for the evolutionary specialization of the digestive system (e.g., four-chambered stomach) of ruminants (Randall et al. 1997; Miller and Harley 2016). Besides birds, teeth reduction or loss is frequently observed in many other tetrapod lineages as well (e.g., toads and turtles) (Davit-Béal et al. 2009; Louchart and Viriot 2011; Barrett 2014). It appears that all the tetrapod lineages with teeth reduction or loss tend to be low-level consumers (e.g., herbivores and insectivores), at least during their early evolution, as exemplified by ancestral birds, and hence may have been faced with relatively high predation pressures. This may imply that high predation pressure might be a common force in facilitating the reduction or loss of teeth. Of course, there are many low-level consumers (e.g., herbivorous dinosaurs) that still retained their teeth. This might be partly attributed to their lineages’ specific low-level predation pressures due to their, for instance, efficient anti-predator strategies (e.g., large body size). In addition to predation pressure, the possible effects of other ecological factors on the evolution of teeth may exist, such as, food types that may determine the reliance of food acquisition and processing on teeth. Most likely, the evolutionary dynamic of teeth may partly reflect a tradeoff between the selections to maintain or strengthen teeth (e.g., acquiring and processing food) and the selections to reduce teeth as a by-product of accelerating feeding to avoid predators.

## Conclusion

Our molecular phyloecological study shows that ancestral birds (e.g., CALB) underwent a dietary shift to be low-level consumers (e.g., fruit, seed, and/or nut eaters), which may have then made them become the prey of potential predators such as gliding maniraptorans (e.g., dromaeosaurids and troodontids), from which ancestral birds descended. Under this predation pressure, the ancestral birds with inherited gliding flight from their immediate gliding maniraptoran predecessors may have then evolved not only powered flight (e.g., flapping flight) as an anti-predator strategy against gliding predatory maniraptorans but also a specialized digestive system as an evolutionary tradeoff of maximizing foraging efficiency and minimizing their exposure to predators (including both gliding and non-gliding predators). Our results suggest that dietary shift-associated predation pressure may have facilitated the evolutionary origin of birds.

## Materials and methods

### Taxa used

We mainly included 95 species in this study. Of the 95 species, 73 species are birds, belonging to 36 orders, representing the majority of living bird orders (36/39) (Gill and Donsker 2018); 22 species are relatives of birds, including five crocodilians, six turtles, and 11 squamates (Fig. 1). For the 73 bird species included, the majority come from Neognathae, with relatively little species of Palaeognathae. For the Palaeognathae species included, the GenBank sequences of many of our focal genes were missing upon our initial sequence analyses, especially for the ostrich (*Struthio camelus*) and the emu (*Dromaius novaehollandiae*), and thus we selected the two species for transcriptome sequencing.

### Sampling, RNA isolation and cDNA library construction

One ostrich (three months old) and one emu (six months old) were used for sampling. The two individuals were the same two individuals used in one of our previous studies (Wu and Wang 2019). And the methods of RNA isolation and cDNA library construction were almost identical to those of that study (Wu and Wang 2019). Briefly, the two active individuals of an artificial breeding company (Quanxin, Daqing) were transported to the laboratory with vegetables and water provided. The two individuals were sacrificed after 24 hours, and an approximately equal amount of tissue from the liver, pancreas, stomach (proventriculus), and small intestine (duodenum) were obtained and mixed. The mixed tissues were preserved in RNA-locker (Sangon Biotech, Shanghai), flash frozen in liquid nitrogen, and then transferred to a −80 °C refrigerator until further processing. The experimental procedures were carried out following an animal ethics approval granted by Northeast Normal University. All experimental procedures in this study were approved by the National Animal Research Authority of Northeast Normal University, China (approval number: NENU-20080416), and the Forestry Bureau of Jilin Province of China (approval number: [2006]178).

We isolated the total RNA of the two samples using TRIzol Reagent (Invitrogen Life Technologies), following the manufacturer’s protocol and instructions. We monitored the RNA degradation and contamination on 1% agarose gels. We checked the RNA purity using the NanoPhotometer^®^ spectrophotometer (IMPLEN, CA, USA). We measured RNA concentration using the Qubit^®^ RNA Assay Kit in a Qubit^®^2.0 Flurometer (Life Technologies, CA, USA). We assessed the RNA integrity using the RNA Nano 6000 Assay Kit of the Agilent Bioanalyzer 2100 system (Agilent Technologies, CA, USA). We constructed the cDNA library using NEBNext^®^Ultra™ RNA Library Prep Kit for Illumina^®^ (NEB, USA) according to the manufacturer’s protocol. Accordingly, mRNA was purified using poly-T oligo-attached magnetic beads. The enriched mRNA was fragmented into small pieces and was then used for the syntheses of the cDNA strands. The cDNA was purified and size-selected using the AMPure XP system (Beckman Coulter, Beverly, MA, USA). Then, PCR was performed, and the PCR products were purified (AMPure XP system). Library quality was assessed on the Agilent Bioanalyzer 2100 system. Paired-ending sequencing was performed using Illumina HiSeq X-ten (Biomarker Technology Co., Beijing).

### Data filtering and de novo assembly

We generated 10.47G bases and 10.45G bases for the ostrich and emu, respectively. We filtered the raw data by removing reads containing adapters, reads containing ploy-N, and low quality reads. Clean reads were assembled using the de novo assembly program Trinity (v2.5.1) (Grabherr et al. 2011) with default parameters. Unigenes were generated, and unigenes longer than 200 bp were retained for subsequent analyses. The transcriptome sequencing data were deposited into the National Center for Biotechnology Information Sequence Read Archive database under accession numbers (SRR12237019-20).

### Genes, sequence abstraction and alignment

The genes annotated in three KEGG digestive system pathways, including carbohydrate digestion and absorption (map04973), protein digestion and absorption (map04974), and fat digestion and absorption (map04975), were used in this study (Fig. 2, Table S5). For these digestive system-related genes, we abstracted their sequences from the studied ostrich and emu. For this, we downloaded the coding sequences of our target genes of *Gallus gallus* from GenBank and used them as query sequences to blast against the unigene pools of the two species using Blastn software. We subsequently annotated these unigene sequences returned by blasting against the NCBI nr/nt database using the online Blastn, and we kept only the unigene sequences with the same gene annotation as that of the query sequences for subsequent analyses. Besides the two species, we also downloaded our target gene sequences from all birds and reptiles with gene sequences available in GenBank (Table S1). For five genes (e.g., *CPA2*, *G6PC*, *PLA2G2E*, *SLC2A5*, and *SLC36A1*), their sequences of the reptile relatives of birds were unavailable, and thus their sequences from mammals and amphibians were used. For our focal genes, those genes (e.g., amylase genes) with sequences unavailable or available for only few bird species were excluded from our analyses, and eventually, 83 genes were retained for subsequent analyses. We aligned gene sequences using webPRANK (http://www.ebi.ac.uk/goldman-srv/webprank/), and individual sequences with long indels and/or lengths that were too short were removed or replaced by other relevant transcript variants. After this pruning, the translated protein sequences of these genes were blasted against the non-redundant protein sequence database to confirm the correctness of the sequence cutting.

### Positive selection analyses

We performed positive selection analyses of genes using branch and branch-site models implemented in the Codeml program of PAML (Yang 2007). For this, an unrooted species tree (Fig. 1) was constructed following published studies (Jønsson and Fjeldså 2006; McKay et al. 2010; Oaks 2011; Guillon et al. 2012; Pyron et al. 2013; Crawford et al. 2015; Wu and Wang 2019) and the Tree of Life Web Project (http://tolweb.org/tree/), with the phylogenetic relationships among bird orders following one genome-level study (Jarvis et al. 2014). We estimated the ratio of non-synonymous to synonymous substitutions per site (dN/dS or ω) and employed likelihood ratio tests (LRT) to determine statistical significance. Positive selection is determined by the value of ω > 1 with a statistical significance.

### Branch model

We performed positive selection analyses of genes along our focal branches using a two-rate branch model. Upon analysis, we labeled our focal branches as foreground branches and the rest were used as background branches. For this model, ω is assumed to be different between foreground branches and background branches, and its goodness of fit was analyzed using the LRT by comparing it with the one-rate branch model that assumes a single ω value across all branches. If a statistically significant value of ω > 1 was detected in a foreground branch, the two-ratio branch model was further compared with the two-ratio branch model with a constraint of ω = 1 of the foreground branch to further determine whether the value of ω > 1 of the foreground branch was supported with statistical significance.

### Branch-site model

We also used a branch-site model (Test 2) to detect positive selection genes for our focal branches. The branch-site model assumes four classes of sites, with site class 0 and site class 1 respectively representing evolutionarily conserved (0 < ω_0_ < 1) and neutral codons (ω_1_ = 1) across branches, and site classes 2a and 2b representing evolutionarily conserved (0 < ω_0_ < 1) and neutral (ω_1_ = 1) codons for background branches, yet allowed to be under positive selection (ω_2_ > 1) for the foreground branches. The goodness of fit of this model was analyzed by using the LRT by comparing a modified model A with a null model with ω = 1 constrained. Positively selected sites were analyzed by an empirical Bayes method.

### Selection intensity analyses

The selection intensity changes of genes were evaluated using RELAX (Wertheim et al. 2015), which is available from the Datamonkey webserver (http://test.datamonkey.org/relax). For the selection intensity analyses, a parameter k value and its statistical significance were estimated given a priori partitioning of test branches and reference branches in a codon-based phylogenetic framework. Intensified selection is indicated by k >1 and is expected to exhibit ω categories away from neutrality (ω = 1), while a relaxed selection is indicated by k < 1 and is expected to exhibit ω categories converging to neutrality (ω = 1). The statistical significance of the k value was evaluated by comparing an alternative model to a null model using LRT, with the former assuming different ω distributions of the test and reference branches, and the latter assuming k = 1 and the same ω distribution of both test and reference branches.

## Acknowledgments

We thank Lin Chen, Yuanqin Zhao, and Li Gu for helping tissue sampling. This research was supported by the National Natural Science Foundation of China (grant number, 31770401) and the Fundamental Research Funds for the Central Universities.

## Author contribution

Y. W. designed research, performed analyses, and wrote the paper.

## Data and materials availability

All data needed to evaluate the conclusions in the paper are present in the paper and/or the supplementary materials.

## Supplementary materials

Tables S1-S5

